# Data-driven analysis identifies female-specific social modulation deficit after chronic social defeat stress

**DOI:** 10.1101/2024.05.08.593167

**Authors:** Heike Schuler, Rand S. Eid, Serena Wu, Yiu-Chung Tse, Vedrana Cvetkovska, Joëlle Lopez, Rosalie Quinn, Delong Zhou, Juliet Meccia, Laurence Dion-Albert, Caroline Menard, Shannon N. Bennett, Catherine J. Peña, Rosemary C. Bagot

## Abstract

**Background:** Chronic social defeat stress is a widely used depression model in male mice. Several proposed adaptations extend this model to females with variable, often marginal effects. We examine the if widely used male-defined metrics of stress are suboptimal in females and reveal sex-specific adaptations.

**Methods:** Using a data-driven method we comprehensively classified social interaction behavior in 761 male and female mice after chronic social witness/defeat stress, examining social modulation of behavioral frequencies and associations with conventional metrics (i.e., social interaction (SI) ratio).

**Results:** Social stress induces distinct behavioral adaptation patterns in males and females. SI ratio leads to underpowered analyses in females with limited utility to differentiate susceptibility/resilience. Data-driven analyses reveal failure of social adaptation in stressed female mice that is captured in attenuated velocity change from no target to target tests (ΔVelocity) and validate this in three female social stress models. Combining SI ratio and ΔVelocity optimally differentiates susceptibility/ resilience in females and this metric reveals resilient-specific adaptation in a resilience-associated neural circuit in female mice.

**Conclusions:** We demonstrate that psychological or physical social defeat stress induces similar deficits in females that is qualitatively distinct from male deficits and inadequately sampled by male-defined metrics. We identify modulation of locomotion as a robust and easily implementable metric for rigorous research in female mice. Overall, our findings highlight the need to critically evaluate sex differences in behavior and implement sex-based considerations in preclinical model design.

## Introduction

Depression affects twice as many women as men (1–4). Despite known sex/gender differences in prevalence, symptom presentation, severity, comorbidities, and treatment response (5–9), research has focused on males. There are growing efforts to include females in preclinical research yet most currently used models and tests for depression-relevant behaviors were developed in males and have not been validated in females (10). Increasing awareness of sex differences in behavior suggests cause for concern (11–16).

In male mice, chronic social defeat stress (CSDS) has yielded important insights into the neurobiology of stress and depression-like states (17–20). CSDS exposes mice to daily bouts of physical defeat followed by sustained psychological stress from sensory exposure to the aggressor across a plexiglass divider. This reliably induces social-avoidance in the majority (∼60%) of male mice (‘susceptible’), and other depression-relevant alterations (17–19). Approximately 40% of male mice (‘resilient’) do not display social avoidance, making this model useful for studying both resilience and susceptibility (19).

Various models aim to extend CSDS to females. Male mice do not readily attack females, necessitating adaptations including chronic witness or vicarious defeat (CSW/DS) with female mice witnessing the physical defeat of a male (21–23) or manipulations to induce direct attacks on females, by masking the female with male sensory stimuli, e.g. male urine or, in non-discriminatory defeat (CSNDS), introducing a male with the female (24,25). Female defeat models generally produce marginal and variable stress effects on social interaction, motivating continued discussion as to how best defeat females. Much effort has focused on developing ‘more robust’ models, assuming female defeat is less robust. The alternative, that defeat induces divergent behavioral adaptations in male and female mice and the conventional male-defined metrics are not optimal for capturing female stress effects, has been largely overlooked.

Recent advances in data-driven analysis and automated behavioral annotation offer a tractable solution by rigorously probing behavior to reveal effects in both sexes. Markerless motion tracking with pose estimation (e.g. DeepLabCut (26–28)) and behavioral classifiers, can comprehensively identify stereotypic behavioral patterns (29–32). Conventional, supervised behavioral classification is constrained by experimenter-defined behaviors of interest, ‘baking in’ untested assumptions. In contrast, unsupervised approaches are less influenced by experimenter and field-wide biases by identifying behavioral motifs without *a priori* conceptual constraints. Here, we employ high-resolution profiling of social stress-induced changes using unsupervised classification to identify novel female-specific stress adaptation. This female-specific social stress signature robustly detects stress effects in standard sized cohorts and is captured by a simple velocity change metric that generalizes to three distinct female defeat models and labs. We demonstrate this metric captures biologically relevant variation in social stress adaptation and identify resilient-specific neural activity patterns. Finally, we provide easily implementable directives for capturing defeat effects in female mice to propel preclinical stress research in females.

## Methods and Materials

### Animals

7-week-old male and female C57BL/6J mice (Jackson, Bar Harbor, ME, USA) were group-housed by sex (n=5/cage) for one week prior to experiments. Retired male CD-1 breeders (approximately 6 months) (Charles River, St Constant, QC, Canada/ Kingston, NY, USA or Raleigh, NC, USA) were single housed. All mice were maintained at 22-25°C, on a 12-h light/dark cycle (lights on at 07:00h), with ad libitum food and water. All procedures were approved by the Animal Care Committee and conformed to McGill University Comparative Medicine and Animal Resources Centre guidelines.

### Chronic social defeat stress

#### CSW/DS

20 cohorts with an average sample size of 38 male and female mice (*n_total_* = 761; *n_Male Ctl_* = 141, *n_Male Stress_* = 216, *n_Female Ctl_* = 174, *n_Female Stress_* = 230) were generated in-house by four experimenters over 4 years. Daily, for 10 days, male mice were placed into the cage of a novel CD-1 aggressor and a female witness placed across the plexiglass divider. After 10 minutes, the female was returned to its home cage (single-housed) and the male moved across the plexiglass barrier until the next daily defeat session.

#### CSNDS

For detailed methods see (33) (n_Female Ctl_ = 5, n_Female Stress_ = 15; n_Cohorts_ = 1). Briefly, daily for 10 days, male and female C57BL/6J pairs were placed in the home cage of a novel Swiss Webster aggressor mouse, with the male mouse place alone for 3 min to prime the aggressor and the female mouse added for an additional 5 minutes. At the end of each defeat session, males were moved across the plexiglass barrier and females were moved to individual cages with aggressor bedding.

#### Urine model

For detailed methods see (*24,34*) (*n_Female Ctl_* = 66, *n_Female Stress_* = 172; *n_Cohorts_* = 8). In short, daily for 10 days female mice were coated in male CD-1 mouse urine prior to being placed in the cage of a CD1 aggressor. After 10 minutes, females were moved across the plexiglass barrier.

### Behavioral testing

Testing occurred during the light cycle under red light and recorded (EthoVisionXT 13, Noldus *or* AnyMaze™ 6.1, Stoelting Co) from top-view for analysis. Mice underwent a two-trial social interaction test (SIT) 24h post CSW/DS (Fig. 1a), and a subset were assessed 24h later in the open field test (OFT).

**Figure 1.**
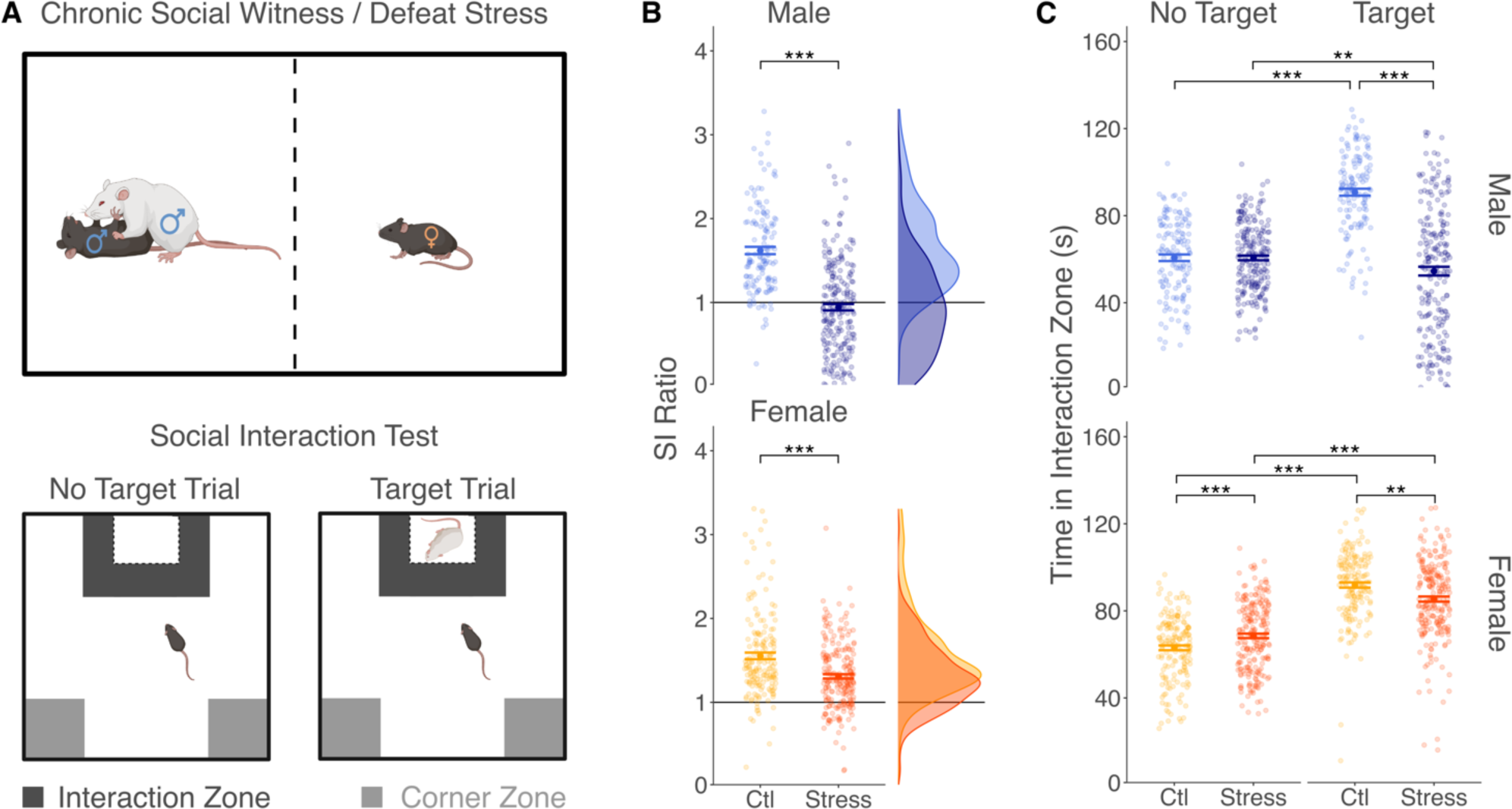
Social interaction ratio is a sensitive metric of stress effects in males but not in females. (A) Schematic illustrates procedure for chronic social defeat and witness defeat and social interaction test. (B) Combined data from 20 cohorts. Social interaction ratio in male (upper panel) and female (lower panel) in control (light color) and stress (dark color) mice represented as individual data points with mean and SEM overlaid (left panel). In a large sample, stress significantly decreases SI ratio in both sexes (ANOVA; <u>Condition:</u> *F_1,738_* = 165.138, *p* < 0.001; <u>Sex:</u> *F_1,738_* = 29.532, *p* < 0.001; <u>Condition x Sex:</u> *F_1,738_* = 35.777, *p* < 0.001; <u>Cohort:</u> *F_19,738_* = 4.309, *p* < 0.001; <u>Sidak Post-Hoc:</u> Control Male *vs* Stress Male: *t-ratio_738_* = 13.324, *p_adj_* < 0.001; Control Female *vs* Stress Female: *t-ratio_738_* = 5.465, *p_adj_ <* 0.001). Notably, there is no sex effect in controls but only in stressed animals (Control Male *vs* Control Female: *t-ratio_738_* = 0.127, *p_adj_ >* 0.999; Stress Male *vs* Stress Female: *t-ratio_738_* = −8.831, *p_adj_* < 0.001). In males, stress markedly shifts the frequency distribution of SI scores whereas in females any shift is minimal (right panel). The variance is larger in stressed females than stressed males, but no difference in variance is observed between sexes in controls (<u>Control:</u> *F_140,173_* = 1.006, *p* = 0.967; <u>Stress:</u> *F_215,229_* = 2.014, *p* < 0.001). The large majority of stressed females remain above 1. (C) Raw time in the interaction zone in No Target and Target trials confirm expected directionality of effect in males (repeated measures ANOVA; <u>Condition:</u> *F_1,338_* = 117.75, *p* < 0.001; <u>Trial type:</u> *F_1,355_* = 32.65, *p* < 0.001; <u>Condition x Trial type:</u> *F_1,355_* = 150.89, *p* < 0.001; <u>Cohort:</u> *F_17,338_* = 117.75; <u>Sidak Post-Hoc:</u> No Target Control *vs* No Target Stress: *t-ratio_675.42_* = −0.067, *p_adj_ =* 0.946; Target Control *vs* Target Stress: *t-ratio_675.42_* =16.073, *p_adj_* < 0.001; No Target Control *vs* Target Control: *t-ratio_355_* = −13.146, *p_adj_* > 0.999; No Target Stress vs Target Stress: *t-ratio_355_* = 3.275, *p_adj_* < 0.01) and minimal yet significant changes in stressed females in both the No Target and Target trial (<u>Condition:</u> *F_1,385_* = 0.199, *p =* 0.655; <u>Trial type:</u> *F_1,402_* = 431.05, *p* < 0.001; <u>Condition x Trial type:</u> *F_1,402_* = 31.04, *p* < 0.001; <u>Cohort:</u> *F_17,385_* = 31.04, *p* < 0.001; <u>Sidak Post-Hoc:</u> No Target Control *vs* No Target Stress: *t-ratio_786.15_* = −4.069, *p_adj_* < 0.001; Target Control *vs* Target Stress: *t-ratio_786.15_* = 3.765, *p_adj_* < 0.01; No Target Control vs Target Control: *t-ratio_402_* = −17.829, *p_adj_* < 0.001; No Target Stress *vs* Target Stress: *t-ratio_402_* = −12.009, *p_adj_* < 0.001). \*\**p_adj_* < 0.01 \*\*\**p_adj_* < 0.001.

### Video analysis

EthoVisionXT 13, Noldus *or* AnyMaze™ 6.1, Stoelting Co automatically quantified velocity and time in interaction zone (IZ; 14 x 24 cm rectangle surrounding mesh enclosure) in SIT to calculate SI ratio (19). Mice with SI ratio > 4 were excluded from analysis (<u>CSW/DS:</u> *n* = 16; <u>Urine model:</u> *n =* 3). Where only distance was quantified (Urine model), velocity was calculated as: distance (cm) / 150 seconds. ΔVelocity was calculated as: velocity (No Target trial) – velocity (Target trial).

A DeepLabCut (DLC; v2.2.1.1 (26–28)) model was trained to track 11 body parts (snout; center, front and back torso; the four extremities; tail base, tail tip and center of the tail). Next, a keypoint-MoSeq (kp-MoSeq) (29) model was trained on 78 No Target and Target SIT trials of control and stressed males and females with the following non-default parameters: *κ* = 1e6; latent dimensions = 5. Snout, center torso, front torso, back torso, tail base and left and right shoulder markers were used to generate repeated behavioral motifs i.e. ‘behavioral syllables’ (29). Frequency of each behavioral syllable was summed per animal/trial. Only syllables present in all cohorts for average > 500 ms (15 frames) per trial were included in analysis. To interpret syllables, we calculated motion, pose and location metrics as described in the supplement.

### Stereotaxic fiber implantation, virus injection and analysis for fiber photometry

Mice were injected with 0.7μl of pGP-AAVrgsyn-jGCaMP7f-WPRE virus (1.85× 10^13^GC/ml; Addgene) in the nucleus accumbens (NAc; A/P: +1.3, M/L: +/-0.60, D/V: −4.9/-4.7), 0.1ul/min for projection-specific GCaMP7f expression. Chronically implantable optic fibers (Neurophotometrics) with 200μm core and 0.37 NA threaded through ceramic ferrules were implanted above infralimbic medial prefrontal cortex (mPFC; A/P: −0.3, M/L: +/-1.90, D/V: −2.80).

To measure calcium-associated fluorescence changes, mPFC to NAc-projecting cells were recorded during the SIT following CSW/DS as previously described ((35); Fig. 7a). Following normalization, No Target and Target trials were concatenated and z-scored for each animal.

### Statistical analysis

Statistical comparisons used (repeated measures) ANOVAs or linear mixed effects models (LME), as appropriate. Specifically, LMEs analyzed syllable data (Fig. 2) and their characteristics (Fig. 4) given nested multilevel data. LMEs analyzed differences in fiber-photometry signal between resilient, susceptible and controls as z-Scores are not generalizable across subjects and analysis employed within-subject comparisons (Fig. 7).

**Figure 2.**
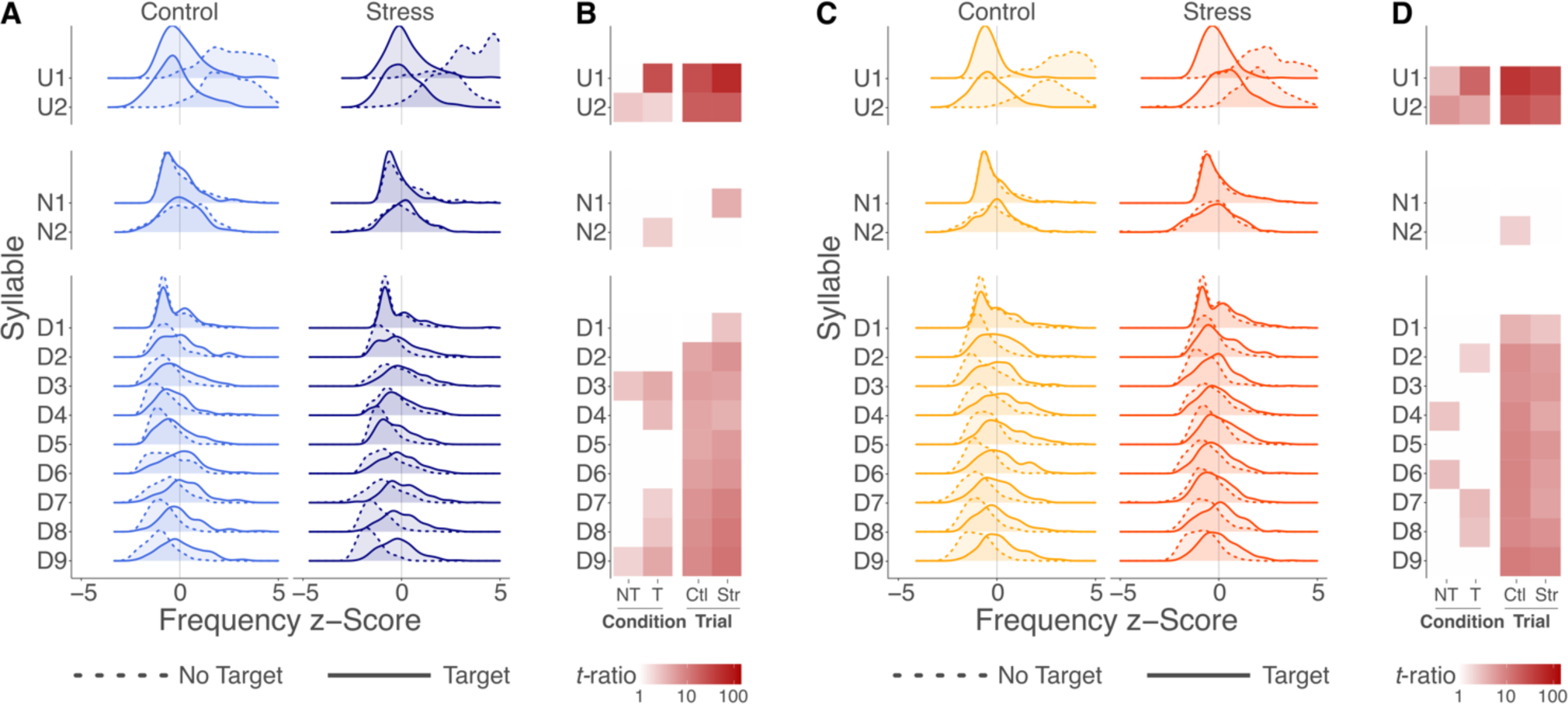
Witness defeat induces modest yet widespread changes across behavioral syllables. Frequency density plots illustrate distribution of z-scored frequency of behavioral syllables in No Target (solid line) and Target (dashed line) trials in (A) male controls (light blue) and stressed mice (dark blue) and (C) female controls (yellow) and stressed mice (orange). Syllables are sorted by social identity of syllables (U, upregulated; D, downregulated; N, no change), the frequency of which is significantly modulated between No Target and Target trials in control mice. Heatmap represents interaction effects of stress condition (Ctr, control; Str, stress) and trial type (NT, No Target; T, Target) on syllable frequency in (B) males and (D) females. Grayscale saturation indicates t-ratio of planned post-hoc contrasts to illustrate each contrast’s effect size, with full statistical data included in Table S3. For each heatmap, the left two columns (**Condition**) compare syllable frequencies between control and stressed animals within No Target (NT, left) and Target (T, right) trials, whereas the right two columns (**Trial**) compare syllable frequencies between No Target and Target trials within controls (Ctl, left) and stressed (Str, right) animals. All non-white squares are *p_adj_* < 0.05.

Principal component analysis (PCA) reduced data dimensionality and identified major sources of behavioral variance in SIT with these factors: syllable frequency differences No Target vs Target, one value/animal/syllable; SI ratio; corner ratio. Male and female models were fit separately, with cohort as a covariate or random factor. Post-hoc tests and planned comparisons were performed using estimated marginal means with Sidak’s multiple comparison correction.

Hedge’s *g* was used as a measure of effect size (36,37). Power calculations were performed with power set to 0.8 and Type-I error (*α*) set to 0.05.

## Results

### SI ratio identifies marginal stress effects in females

We first examined effects of CSW/DS on social interaction ratio, the standard measure of stress effects in male defeat mice. CSW/DS reduced SI ratio in both males and females compared to non-stressed controls (Fig. 1b), however this effect is substantially more robust in males, with an effect size (*t_313.49_* = 11.69; *Hedge’s g* = 1.25) more than double that in females (*t_311.11_* = 5.19; *Hedge’s g* = 0.54). While this very large sample (*n_Total_* = 761; see methods) revealed a statistically significant effect in females, a standard sample of 8 to 15 animals would yield only 0.17 to 0.3 probability of detecting this effect. That is, 55 females per group are needed to achieve adequate power comparable to 12 males per group. Notably, this effect size difference is driven by sex differences between stressed but not control animals (Fig. 1b). Further, the variance is higher in male than female stressed, but not control mice (Fig. 1b). Additionally, a much smaller proportion of females than males are classified as susceptible (<u>Males:</u> 57%; <u>Females:</u> 22%), further constraining the utility of the model in females.

We then examined raw time in IZ, from which SI ratio is calculated. In males, we confirmed that controls spend more time in IZ during the Target than the No Target trial, and defeated males show the opposite (Fig. 1c), with no difference between control and stressed males in the No Target trial. In contrast, female effects were not specific to the Target trial (Fig. 1c). While both controls and witnesses spend more time in IZ when a target is present than without a target, compared to controls, witnesses spend less time in IZ in the Target trial, but more time in the IZ in the No target trial. This indicates females are not less affected by stress but that the stress effect is distinct from males.

### Unsupervised behavioral classification reveals widespread effects in female mice

The modest effects on SI ratio in females could indicate limited efficacy of CSW/DS in females, or alternatively suboptimal sensitivity of SI ratio, a male-defined metric, in capturing stress effects. To assess this, we performed unsupervised behavioral classification of the SIT, including 13 behavioral syllables generated by kp-MoSeq in the analysis. As raw frequencies differed strongly between syllables (Fig. S1) we z-scored syllable frequency to avoid high-frequency syllables overpowering the analysis. To identify social behavior syllables, we grouped syllables according to modulation of syllable frequency between non-social (No Target) and social (Target) trials in control mice (Table S1): Upregulated syllables (U: Upregulated) are more frequent and downregulated syllables (D: Downregulated) are less frequent during Target than No Target trials, with constant frequency of no change syllables (N: No Change) across trial types. CSW/DS does not alter the social identity of behavioral syllables: Upregulated syllables remain upregulated and downregulated syllables remain downregulated in both defeated males and witness females (Table S2, Fig. 2a-d). We then asked how stress, trial-type and sex might modulate syllable frequency. Predicting syllable frequency from syllable identity, condition, and trial type identified several interaction effects in both sexes (Table S2, Fig. 2b,d) with mostly modest sized effects. This suggests that CSW/DS induces widespread yet modest changes across social syllables, rather than altering the frequency of a few select syllables.

### Dimensionality reduction identifies distinct sources of stress-related variance in males and females

Having observed widespread yet individually modest syllable changes, we used dimensionality reduction to distill and quantify global behavioral regulation patterns. We performed a principal component analysis (PCA) on SI ratio, corner ratio, and syllable trial-type frequency difference combined across sex (male, female) and condition (control, stress). Visualizing the first two principal components reveals that, while control and defeated males largely separate along PC1, control and witness females separate predominantly along PC2 (Fig. 3a). Factor loadings reveal that PC1 is primarily driven by SI and Corner ratio (Fig. 3b), consistent with the large effect size for standard location-based metrics in male mice. In contrast, PC2 is driven by syllable trial-type frequency differences. We further observed that syllables segregate by social identity with Upregulated syllables loading positively and Downregulated syllables loading negatively onto PC2 (Fig. 3b). This can be explained by attenuated social modulation of behavior in stressed females (Fig. 3c). That is, in the presence of a social target, both increasing upregulated syllables and decreasing Downregulated syllables are attenuated in stressed females compared to controls. We summarized this observation in a ‘Social Modulation Score’ (SMS; Fig. 3d), calculated as the average absolute difference between No Target and Target trial syllable frequencies. In contrast, in stressed males increasing Upregulated syllables in the presence of a social target is more pronounced relative to controls, with no impact on No Change or Downregulated syllables. Modulation of Upregulated syllables in stressed males is driven largely by single syllable, U1, and closely related to SI ratio (Fig. S2).

**Figure 3.**
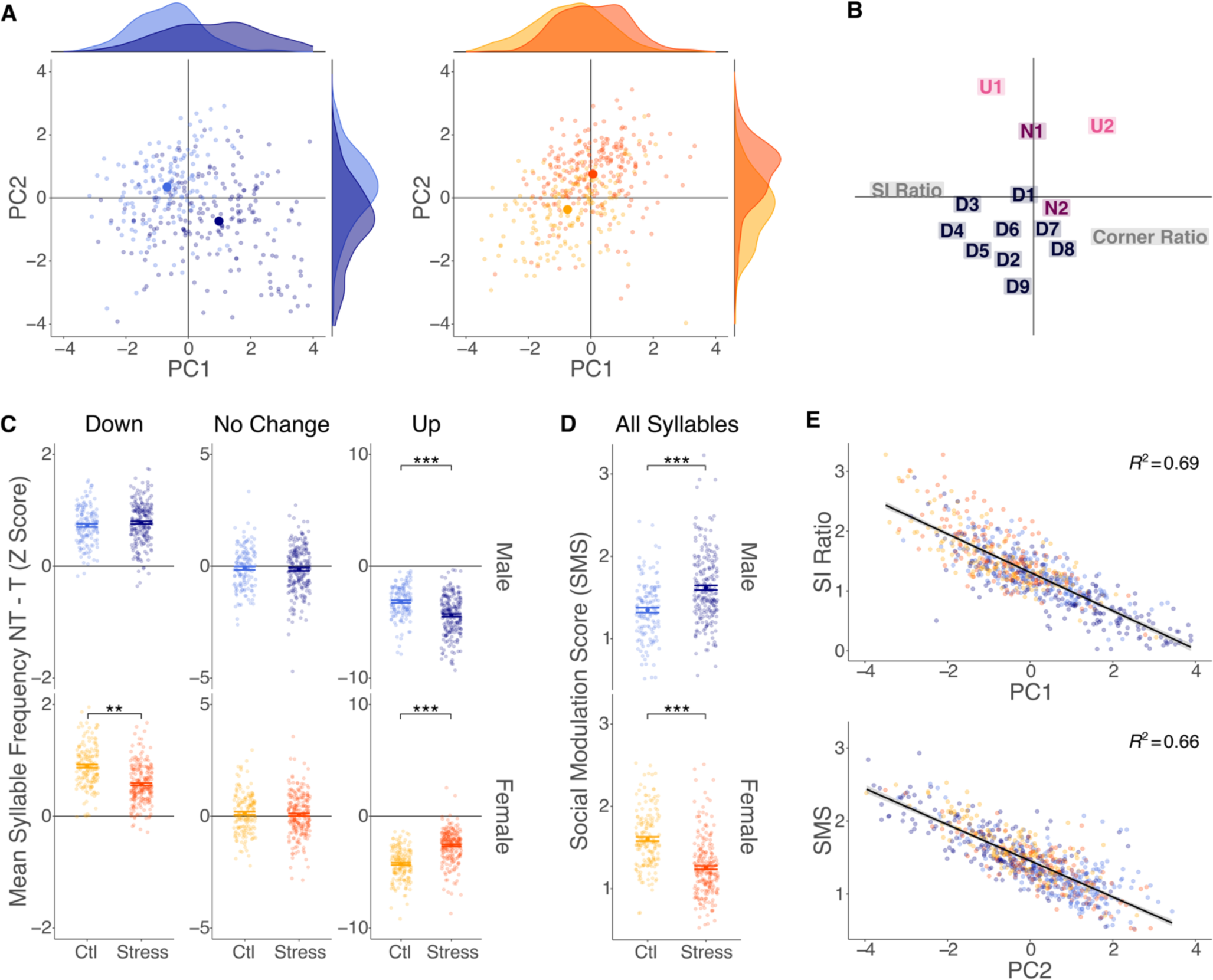
Principal component (PC) analysis of syllable frequency change across trial type and social interaction and corner ratios reveals that stress-associated variance loads onto orthogonal axes in males and females. (A) The combined principal components analysis of males and females represented segregated by sex for clarity. PC1 differentiates male stress phenotype (left panel) whereas PC2 differentiates female stress phenotype (right panel) represented as both individual points and frequency distribution of PC scores. (B) Standard metrics of SI and corner ratio primarily load on PC1 whereas changes in social syllable frequency primarily load on PC2. Color scale indicates degree of syllable social modulation. (C) Average z-scored frequency difference of syllables shows distinct patterns of syllable modulation in males (upper panel) and females (lower panel). In males, stress-induced changes are restricted to upregulated syllables (rmANOVA; <u>Condition:</u> *F_1,338_* = 58.18, *p* < 0.001; <u>Social identity:</u> *F_2,710_* = 1398.33, *p* < 0.001; <u>Condition x</u> <u>Social identity:</u> *F_2,710_* = 28.67, *p* < 0.001; <u>Cohort:</u> *F_17,338_* = 3.58, *p* < 0.001; <u>Sidak Post-Hoc:</u> Control U *vs* Stress U: *t-ratio* = 10.055, *p_adj_* < 0.001; Control N *vs* Stress N: *t-ratio* = 0.3, *p_adj_* = 0.764; Control D *vs* Stress D: *t-ratio* = −0.484, *p_adj_* = 0.628). In females, both up and downregulation is attenuated (<u>Condition:</u> *F_1,385_* = 92.87, *p* < 0.001; <u>Social identity:</u> *F_2,804_* = 1623.14, *p* < 0.001; <u>Condition x Social identity:</u> *F_2,804_* = 96.08, *p* < 0.001; <u>Cohort:</u> *F_17,385_* = 5.271, *p* < 0.001; <u>Sidak Post-Hoc:</u> Control U *vs* Stress U: *t-ratio* = −16.681, *p_adj_ <* 0.001; Control N *vs* Stress N: *t-ratio* = 0.028, *p_adj_* = 0.978; Control D *vs* Stress D: *t-ratio* = 3.143, *p_adj_* < 0.01). (D) This is confirmed by the average absolute change across all syllables i.e. the ‘social modulation score’ (SMS) (ANOVA; Males: <u>Condition:</u> *F_1,338_* = 45.518, *p* < 0.001; <u>Cohort:</u> *F_17,338_* = 3.823, *p* < 0.001; Females: <u>Condition:</u> *F_1,385_* = 115.852, *p* < 0.001; <u>Cohort:</u> *F_17,385_* = 4.479, *p* < 0.001). (E) Correlational analyses confirm that PC1 strongly correlates with SI ratio (*r_Pearson_* = 0.83, *R^2^* = 0.69) and PC2 strongly correlates with the SMS (*r_Pearson_* = 0.81, *R^2^* = 0.66). \**p_adj_* < 0.05 \*\**p_adj_* < 0.01 \*\*\**p_adj_* < 0.001.

This data-driven approach confirms the behavioral impact of CSW/DS in female mice yet, this impact is qualitatively distinct from effects in males and so a single criterion will not optimally capture stress effects in both sexes. PC1 separates control and stressed male and strongly correlates with SI ratio (Fig. 3e), independently validating this metric for use in males. In contrast, control and stressed females segregate primarily on PC2, which strongly correlates with SMS (Fig. 3e), indicating more widespread behavioral alteration not captured by conventional location based metrics, such as time in IZ or corners.

### Velocity is a unifying feature and heuristic for social modulation

A major advantage of SI ratio is that it can be easily calculated with video without advanced analysis. We asked if a simple metric might exist for female mice. Stress induced opposing effects on Up- and Downregulated syllables that were nevertheless concordantly regulated within social identity syllables. This suggested a potential kinematic/location-based motif unifying Up-or Downregulated syllables while also distinguishing them. To identify the key features distinguishing Up- and Downregulated syllables, we used a restricted set of kinematic features (motion, pose, and location metrics) to describe and compare Up- and Downregulated syllables. We found that Up- and Downregulated syllables differ in motion and pose metrics, with Downregulated syllables defined by increased motion and longer pose (Fig. 4a consistent with increased velocity and distance. When mice move at higher speed, head and spine length increase in a more extended position. Location metrics account for less variability between Up- and Downregulated syllables (Fig.4a). This suggests that attenuated social modulation in female witness mice is driven by increased usage of high-locomotion syllables and decreased usage of low-locomotion syllables when the social target is present.

**Figure 4.**
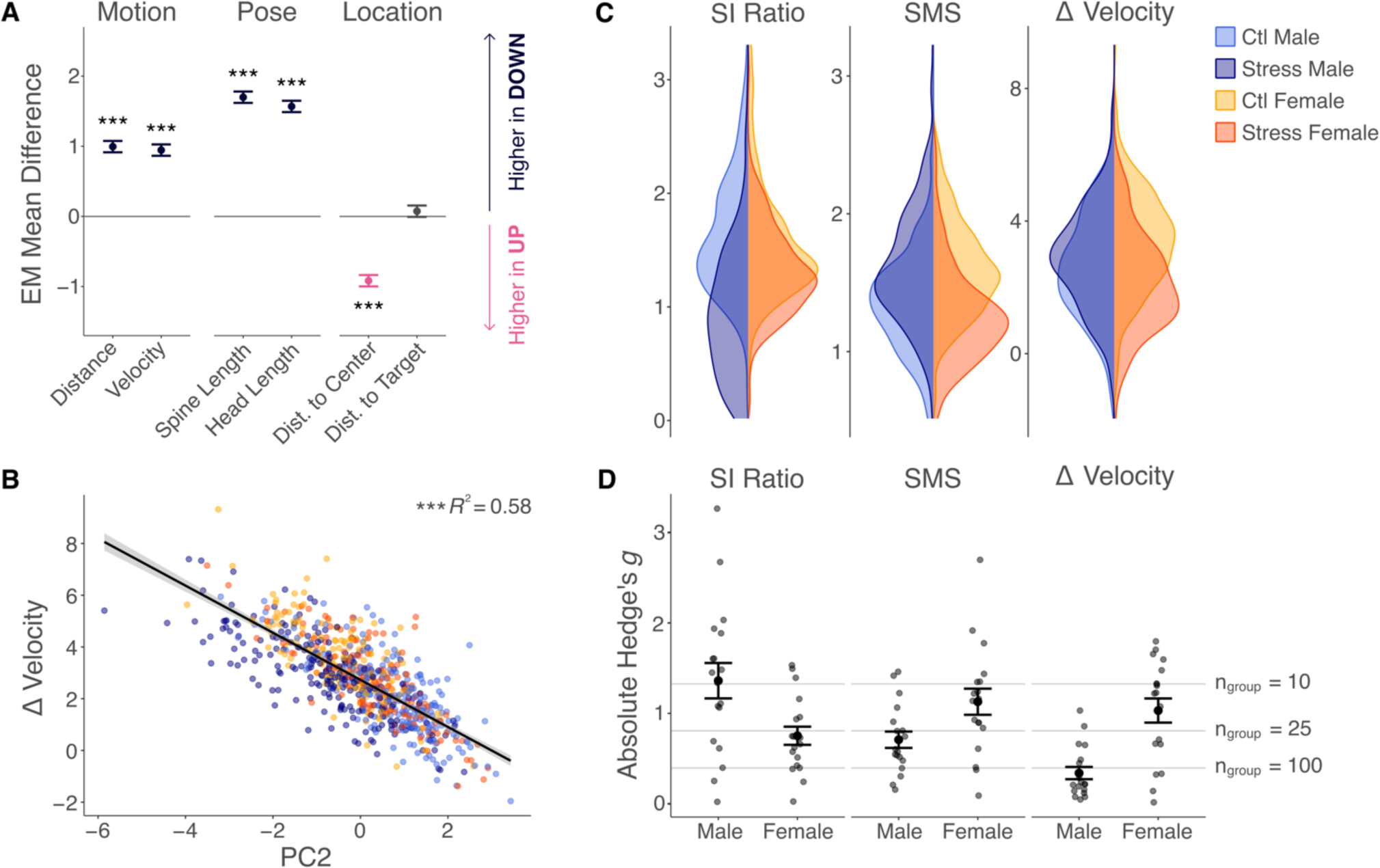
Investigation of syllable features identifies velocity as a unifying feature and a heuristic for quantifying female, but not male, stress phenotype. (A) Contrasts between up- and down-regulated syllables’ estimated marginal means of motion (<u>Sidak Contrast:</u> Velocity U *vs* D: *t-ratio_8360_* = 22.897, *p <* 0.001; Distance U *vs* D: *t-ratio_11837_* = 24.1, *p <* 0.001), pose (<u>Sidak Contrast:</u> Head Length U *vs* D: *t-ratio_8360_* = 37.978, *p <* 0.001; Spine Length U *vs* D: *t-ratio_8360_* = 41.159, *p <* 0.001) and location (<u>Sidak Contrast:</u> Dist. to Target U *vs* D: *t-ratio_8360_* = 1.805, *p <* 0.05; Dist. to Center U *vs* D: *t-ratio_8360_* = −22.179, *p <* 0.001) characteristics demonstrate that downregulated syllables are high-velocity, long-distance syllables (motion) marked by extended head and spine length (pose). The full statistical model is reported in Table S3. (B) ΔVelocity correlates strongly with PC2 (*r_Pearson_* = −0.76, *R^2^* = 0.58, *p* < 0.001) and the SMS (not shown; *r_Pearson_* = 0.7, *R^2^* = 0.49, *p* < 0.001), validating its utility as a heuristic replacing SMS. (C) Visualization of sex and condition-specific distributions and (D) absolute effect size estimates for individual cohorts verify that SI ratio serves as a useful metric to capture the stress effect in males, whereas both SMS and ΔVelocity pick up a stress effect in females.

We found that motion characteristics differentiate Up- and Downregulated syllables which define the SMS to distinguish witness and control females. We reasoned that a single motion characteristic, such as velocity, may provide a heuristic measure as a simple alternative to the SMS. Velocity is a standard output calculated by both commercial and open source behavioral tracking software (e.g., AnyMaze, EthoVision, ezTrack; (38)), so would be easily implementable in any lab already quantifying stress effects in defeat models. Indeed, change in velocity (ΔVelocity) from No Target to Target is highly correlated with both PC2 and the SMS (Fig. 4b). This is driven largely by a higher velocity in stressed than in control females during the Target trial and it is not observed in males (Fig. S3).

We then compared performance of ΔVelocity to both SI ratio and SMS. Visualizing distributions for SI ratio, SMS and ΔVelocity in control and stressed males and females (Fig. 4c) highlights sex differences in behavioral metrics: Compared to controls, SI ratio is downregulated in stressed males but not females, whereas both SMS and ΔVelocity are downregulated in stressed females but not males. Effect size calculations confirm little advantage of using SMS over ΔVelocity. Both SMS and ΔVelocity substantially increase effect size estimates for females compared to SI ratio. Both metrics require 14-16 animals per group for adequate power (*g_SMS_* = 1.13, *g_ΔVelocity_* = 1.03; Fig. 4d).

### Attenuated ΔVelocity generalizes across female social stress models

CSW/DS attenuated ΔVelocity in females but not males, and SI ratio was strongly modulated in males but not females. We hypothesised this reflects a sex difference in adaptation to social stress. Alternatively, it the *type* of stress may dictate behavioral adaptation: Female mice experienced a psychological, vicarious defeat whereas male mice a direct physical defeat. We, therefore examined the generalizability of our findings to female defeat models using direct attacks to determine if the observed phenotype is shaped by the nature of the chronic stressor or reflects a fundamental sex difference.

We examined chronic non-discriminatory defeat stress (CSNDS), wherein females experience limited physical attacks from proximity to male defeat mice and the urine model, wherein female mice experience substantial physical attacks as they are disguised in male urine. In both models and similar to CSW/DS, defeat attenuated ΔVelocity, although in the urine model, this did not reach statistical significance (Fig. 5b). SI ratio is not significantly decreased in CSNDS stressed females (Fig. 5c) and in the urine model SI ratio is reduced with an effect size similar to that observed in CSW/DS (Fig. 5c). This indicates that direct physical attacks did not increase SI ratio sensitivity for capturing stress effects in female mice, confirming that the metric, not the model, is suboptimal.

**Figure 5.**
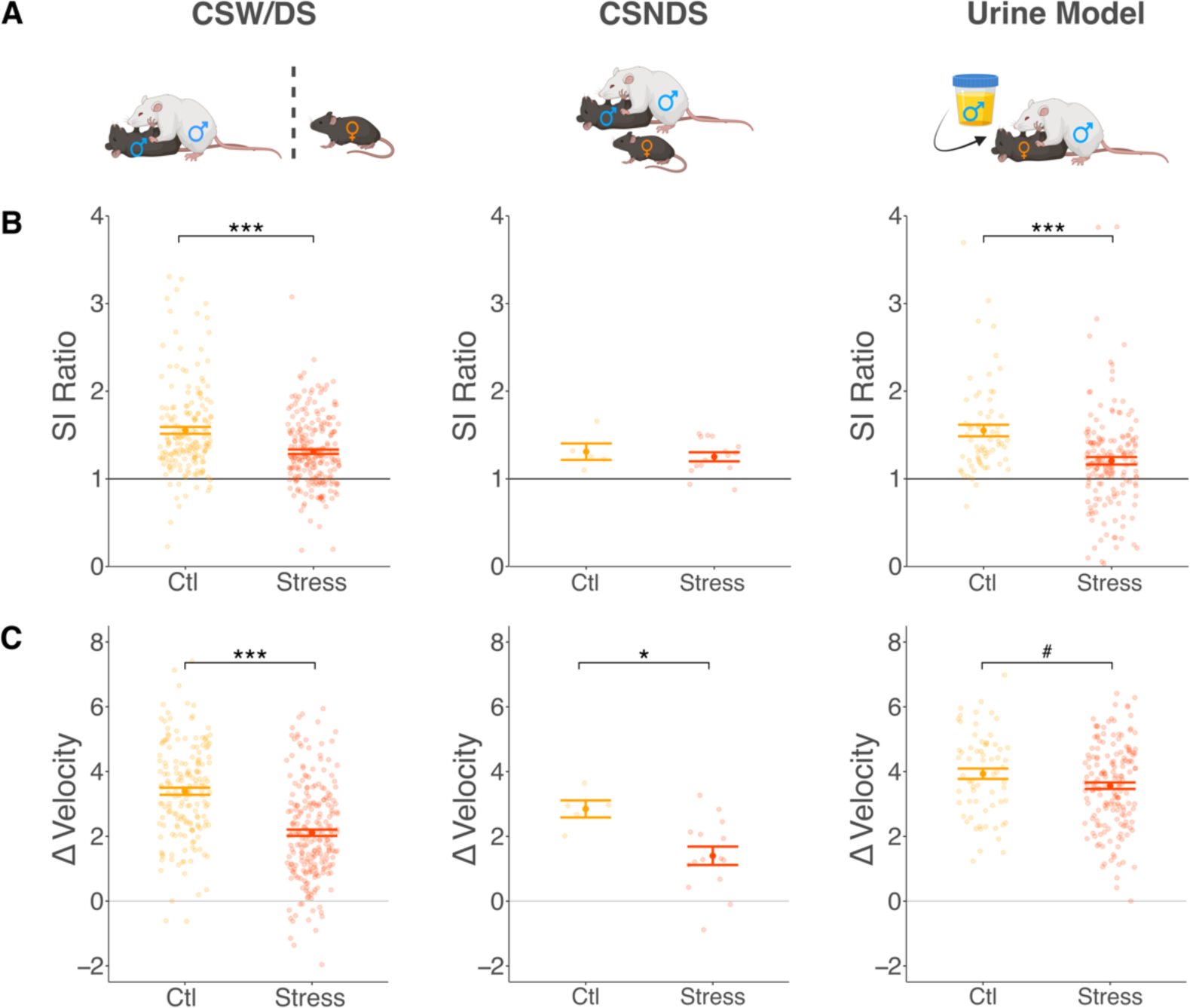
ΔVelocity is decreased after vicarious and physical forms of chronic stress. (A) Different female models of chronic stress expose animals to emotional/vicarious stress (CSW/DS), a low number of attacks by proximity (CSNDS), or to physical attacks similar to those experienced by males in CSDS (Urine defeat model). (B) SI ratio is similarly decreased after both CSW/DS (ANOVA; <u>Condition:</u> *F_1,385_* = 37.972, *p <* 0.001, <u>Cohort:</u> *F_17,385_* = 8.277, *p <* 0.001; *Hedge’s g* = 0.54) and Urine model defeat (<u>Condition:</u> *F_1,226_* = 19.386, *p* < 0.001, <u>Cohort:</u> *F_7,226_* = 2.612, *p* < 0.05; *Hedge’s g* = 0.63), and not significantly altered after CSNDS (*F_1,18_* = 0.318, *p* = 0.58; *Hedge’s g =* 0.28). (C) ΔVelocity is significantly decreased after both CSW/DS (<u>Condition:</u> *F_1,385_* = 97.360, *p* < 0.001; <u>Cohort:</u> *F_17,385_* = 7.426, *p* < 0.001) and CSNDS (*F_1,18_* = 7.693, *p* < 0.05), with a trending decrease after Urine model defeat (<u>Condition:</u> *F_1,226_* = 3.751, *p* = 0.054; <u>Cohort:</u> *F_7,226_* = 3.892, *p* < 0.001). ^#^*p* = 0.054 \**p* < 0.05 \*\*\**p* < 0.001.

### Combining ΔVelocity and SI Ratio optimally differentiates resilient and susceptible females

In male mice, SI ratio differentiates resilient and susceptible phenotypes using a cut-off of 1 with values <1 indicating social avoidance and >1, social engagement (19). This corresponds to the distribution in control mice, wherein the large majority have SI ratio >1. To identify a resilience/susceptibility threshold for ΔVelocity, we used a distribution-based metric. Examining SI ratio and ΔVelocity together reveals an inverted U-shape relationship consistently across female stress models: SI values < 1 are negatively related to ΔVelocity, whereas SI values >1 are positively related to ΔVelocity (Fig. 6a). This is not observed in males (Fig. 6a). To determine susceptibility, we propose a ΔVelocity cut-off of 1.5 to approximate the one-sided 10% z-confidence interval of the CSW/DS female control population, thereby designating stressed animals with ΔVelocity values outside the commonly observed range in control females. Given that SI ratio is also modestly regulated in females in addition to the much larger effect on ΔVelocity, we propose additionally designating as susceptible females with SI ratio less than 1, irrespective of ΔVelocity, to account for the bimodal relationship between the two metrics.

**Figure 6.**
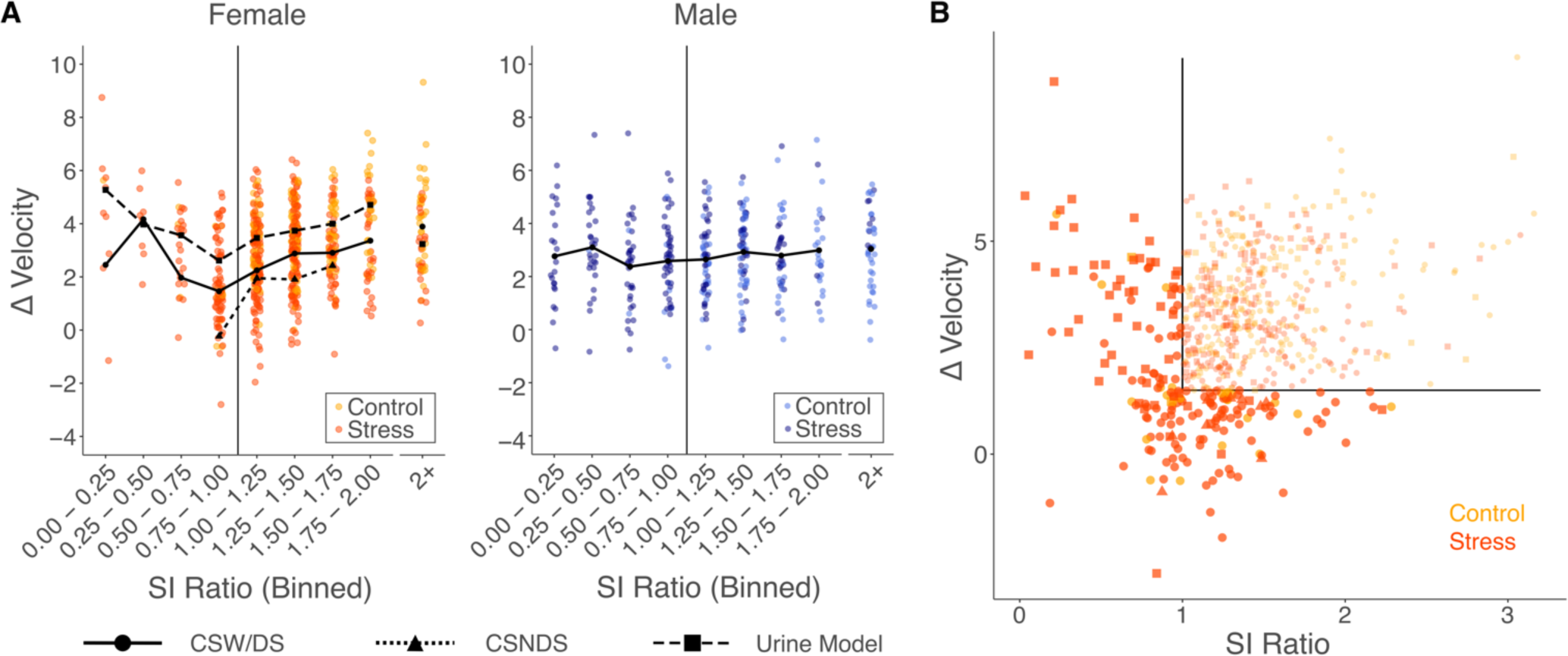
Susceptibility and resilience in female mice should be defined using both SI ratio and ΔVelocity. (A) SI ratio and ΔVelocity have a bimodal relationship in females (left panel) but not in males (right panel). SI ratio < 1 is negatively associated with ΔVelocity, whereas SI ratio > 1 is positively associated. SI ratio was binned to allow for better visualization in the context of the sparsity of low SI ratio values. (B) Using a dual cut-off identifies stressed females that fall outside the control distribution. We propose to retain the cut-off of 1 for SI ratio and suggest classifying females with ΔVelocity < 1.5 as susceptible based on the control population distribution (*mean* = 3.425; *SD* = 1.516; *Conf. Interval* = [1.485, ∞]).

In male mice, in addition to susceptible-specific deficits in social interaction, defeat induces an anxiety-like phenotype in both susceptible and resilient animals, most commonly observed is reduced open field centre exploration. We did not observe clear modulation of centre exploration in female mice, however stressed females showed increased velocity in OFT (Fig. S4). As in males, the increased velocity is common to resilient and susceptible females suggesting a similar dissociation of social stress effects on social reward and anxiety-like behavior, and further supporting the specificity of ΔVelocity to social modulation in the SIT.

### Social modulation of mPFC-NAc neural activity in resilient, but not susceptible, females

Using a data-driven approach we identified novel social stress induced behavioral variability uniquely in female mice. A major strength of male CSDS is in revealing distinct neurobiological mechanisms of susceptibility and resilience. We tested the utility of the novel ΔVelocity+SI metric in identifying biological mechanisms of resilience and susceptibility in females. The NAc and mPFC are implicated in differential adaptation to chronic stress in both sexes (18,34,39,40) and in male mice, increased mPFC-NAc circuit activity is pro-resilient (41). We hypothesised that mPFC-NAc neural activity would also distinguish resilient and susceptible females.

To probe mPFC-NAc activity, we injected retrograding AAV-GCaMP7f in NAc and recorded Ca^2+^-associated fluorescence from optic fibers in mPFC during a SIT following CSW/DS. We compared mean Z-scored activity during the No Target and Target test in control mice and mice identified as susceptible or resilient by the ΔVelocity+SI metric. We observe that, mPFC-NAc neural activity in resilient mice is modulated by target presence, increasing during the Target trial arena-wide (Fig. 7b) and in IZ (Fig. 7c). Susceptible females fail to show this social modulation, consistent with impaired social modulation of behavior. This validates the utility of this novel metric for capturing biologically meaningful variation in stress adaptation. Applying the combined ΔVelocity+SI ratio cut-off identified five of the fourteen defeated mice as susceptible and nine resilient whereas the standard SI metric would have misclassified three of these five susceptible mice as resilient.

**Figure 7.**
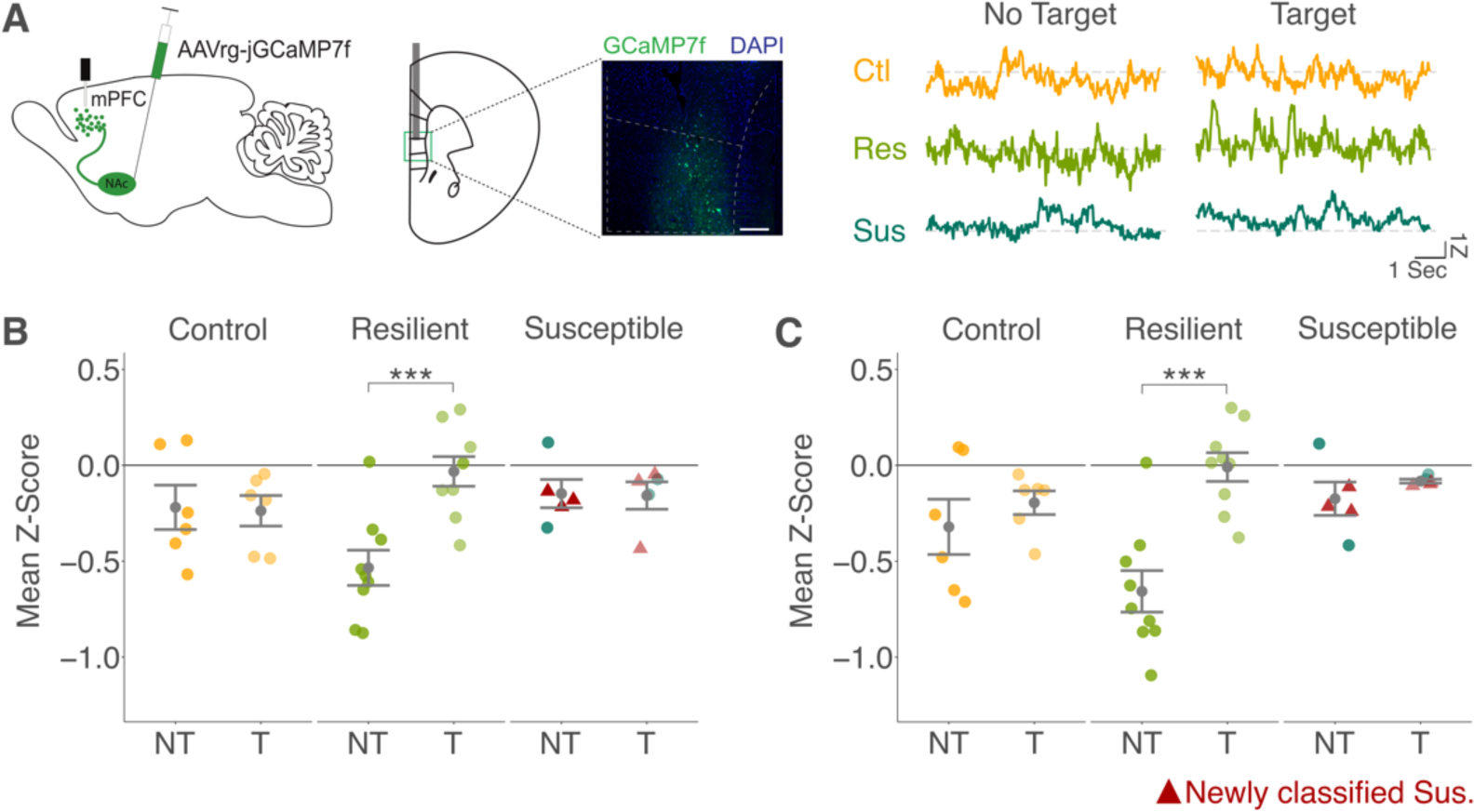
mPFC to NAc pathway responses differ between resilient and susceptible females during SIT. (A) Animals were implanted with Z-scored representative traces of one animal per group are shown in the right panel during the No Target and the Target trial. Mean Z-Score in (B) the whole arena and (C) in the interaction zone show trial-type discrimination in resilient mice (<u>Sidak Contrasts NT *vs* T:</u> Arena: *t-ratio_17_* = −4.583, *p_adj_ <* 0.001; IZ: *t-ratio_17_* = −5.483, *p_adj_ <* 0.001), but not in susceptible (<u>Sidak Contrasts NT *vs* T:</u> Arena: *t-ratio_17_* = 0.069, *p_adj_ =* 0.946; IZ: *t-ratio_17_* = −0.578, *p_adj_ =* 0.571) or control females (<u>Sidak Contrasts NT *vs* T:</u> Arena: *t-ratio_17_* = 0.134, *p_adj_ =* 0.894; IZ: *t-ratio_17_* = −0.869, *p_adj_ =* 0.397). Animals that would have been classified as resilient based on SI ratio alone are displayed as triangle shaped red points. Full model results are reported in Table S4. \*\*\**p_adj_* < 0.001.

## Discussion

Uncovering depression mechanisms in females requires robust preclinical models yet female social defeat models have not been widely adopted, with reports of marginal efficacy. We demonstrate the apparent lack of efficacy is largely attributable to limited generalizability of male-defined behavioral metrics. We show that the conventional metric, SI ratio, is underpowered to detect stress effects in females. To discover relevant female behavioral variation, we applied unsupervised behavioral classification, revealing widespread, yet modest changes characterized by lack of social modulation distinct from social target proximity effects in males. Both a female-optimized SMS and a simple velocity-based heuristic sensitively capture the female phenotype across three defeat variants. This demonstrates the efficacy of female defeat models, and that male-defined behavioral metrics are suboptimal for detecting female behavioral adaptation. Beyond developing an essential research tool, this emphasizes the need to consider sex in all aspects of experimental design.

There is growing recognition of sex differences in behavior and stress adaptation (11–16). We demonstrate that defeat stress discretely changes male behavior, primarily altering location-based SIT metrics captured by the SI ratio. Conversely, in females, stress attenuates social modulation of a large behavioral repertoire, and this is captured by the SMS. This female-specific stress-induced phenotype is driven by increased high-locomotion and decreased low-locomotion behavioral syllables in the presence of a social target. This observation led to discovering that a simple velocity change (ΔVelocity) metric efficiently captures this variation, bypassing the need for pose estimation and behavioral classifiers and minimizing required sample sizes. To further limit numbers of animals required for sufficiently powered experiments, we suggest adopting published methods to leverage historical control data to optimize feasibility (42). Using a population-based cut-off to define susceptible and resilient female mice reveals that variable social modulation of velocity identifies differential adaptation in a resilience-associated neural circuit. Notably, we also observed increased velocity in all stressed females in OFT, but not in males, suggesting a general hyperlocomotion in addition to susceptible-specific deficits in social modulation. This female-specific hyperlocomotion phenotype is reminiscent of psychomotor symptoms in depression and other stress-related psychiatric disorders, wherein evidence exists of gender differences (43,44).

ΔVelocity is a broadly useful and applicable metric, detecting robust stress effects in data across multiple laboratories and female defeat variants, including witness defeat, non-discriminatory defeat, and urine models. Having developed a sensitive, robust metric in females, we demonstrated that both physical and/or psychological defeat models induce similar stress phenotypes. This challenges the notion that “better” female models are needed, or that direct physical attack is necessary, while underscoring the generalizability of the metric.

This work does not invalidate previous findings with female defeat. However, in studies relying on SI ratio, a subset of susceptible females may have been wrongly classified as resilient, obscuring differences. An integrated ΔVelocity+SI cut-off more accurately classifies susceptible females, facilitating novel biological insight into resilience and susceptibility. Applying this metric we reveal aberrant social modulation in mPFC-NAc activity in susceptible mice, mirroring behavioral deficits and consistent with findings implicating this pathway in resilience in males (41).

Overall, our findings illustrate the critical importance of rigorously assessing male-defined behavioral metrics to test utility in females. Harnessing advances in data-driven analysis to rigorously examine female behavior, we revealed previously unrecognized sex differences in behavioral adaptation to social stress. Applying this insight, we defined an easily implementable metric validated in three female defeat models. This will enable robust preclinical depression research in female mice to advance mechanistic understanding of stress susceptibility and depression.

## Acknowledgments

All data will be made publicly available upon publication here: https://osf.io/g3brw/. All code used to statistically analyze and plot the data can be found here: https://github.com/heike-s/FemaleStressPhenotype/. Graphics in Figs. 1 and 5 were created with BioRender.

Funding sources: CIHR, Ludmer Centre for Neuroinformatics and Mental Health, FRQS.

