## Supplements File for "Data-driven analysis identifies female-specific social modulation deficit after chronic social defeat stress"

### Supplemental Materials & Methods

*Chronic social witness defeat stress*

Male CD-1 mice were screened for aggressive behavior, meeting criteria for inclusion as aggressors if displaying ≥5 attacks/5 minutes toward a novel C57BL/6J male intruder across 3 days of screening, with an attack latency of <30 seconds on the final day of screening. The day before starting chronic social witness defeat stress (CSW/DS), CD-1 aggressor mice were individually housed on one side of a 46 x 24 cm cage, divided in half by a clear perforated plexiglass divider. On each of the 10 days of defeat, male C57BL/6J mice were placed into the cage of a novel CD-1 aggressor mouse for 10 minutes. A female C57BL/6J mouse was placed into the compartment across the plexiglass divider to witness the defeat. At the end of each daily defeat bout, the female mouse was returned to its home cage (single-housed) and the male C57BL/6J was moved across the plexiglass barrier where it remained until the start of the next daily defeat bout. This provided for continued sensory exposure to the CD-1 aggressor in male mice as per the standard chronic social defeat protocol (1,2). This was repeated daily with the male and female mouse exposed to a novel CD-1 aggressor each day. Control male and female C57BL/6J were pair-housed by sex across a perforated plexiglass divider in a 46 x 24 cm cage and weighed daily, but not otherwise handled, and remained in their assigned cage for the duration of the experiment. On the final day of defeat, all mice were individually housed in standard mouse cages before starting behavioral testing the next day.

*Validation models*

**Non-discriminatory social defeat.** Detailed methods are reported in (3). In brief, mice experienced chronic non-discriminatory social defeat stress, a model for depression and social stress-related disorders. In this paradigm, female and male C57BL/6J pairs were simultaneously placed into the home cage of a novel Swiss Webster aggressor mouse. An experimental male mouse was introduced to an aggressor's cage first for ~3 minutes, followed by an experimental female mouse for an additional 5 minutes. Male mice were then moved across a perforated plexiglass barrier within the aggressor cage for the remainder of the day, for sensory stress without physical interaction. Female mice were removed to individual cages with aggressor bedding for the remainder of the day. Experimental mice were introduced to a new aggressor each day for 10 days, and male and female mice rotated in different directions so that trios were unique each day. While aggressors were observed to attack female mice, attacks were predominantly directed at male mice. No copulations were observed. Control mice were housed in a standard mouse cage in pairs of the same sex, separated by a perforated plexiglass barrier. At the conclusion of social defeat, all mice were rehoused in clean cages.

**Urine model.** Detailed methods are reported in (4,5). Briefly, CSDS using urine was performed as previously described by Harris et al. (5). CD-1 mice were screened for aggressive behavior during inter-male social interactions for three consecutive days based on previously described criteria and housed in the social defeat cage (26.7 cm width × 48.3 cm depth × 15.2 cm height, Allentown Inc) 24 h before the start of defeats (day 0) on one side of a clear perforated Plexiglas divider (0.6 cm × 45.7 cm × 15.2 cm, Nationwide Plastics). Each female mouse was paired with the urine of a particular male CD-1 mouse throughout the entire course of CSDS. Each day, urine was applied to the base of the tail (20 μL), vaginal orifice (20 μL) and upper back (20 μL) of the female mouse then it was immediately subjected to physical interactions with an unfamiliar CD-1 AGG for 10 min. After antagonistic interactions, experimental mice were removed and housed on the opposite side of the social defeat cage divider, allowing sensory contact, over the subsequent 24 h period. Throughout the sessions, mice were monitored for aggressive interactions and mounting behaviors. A session was immediately stopped if persistent mounting or fighting causing physical wounding occurred. Unstressed control mice were housed two per cage on either side of a perforated divider and rotated daily in a similar manner without being exposed to the CD-1 AGG mice. Experimental and control mice were singly housed after the last bout of physical interaction and the social interaction (SI) test was conducted 24 h later. Physical wounding was scored at the time of tissue collection, 24 h after the SI test, and consisted of counting the number of tail bites on the experimental animals as well as the surface area ($cm^2$) of lower back lacerations, if applicable.

*Behavioral testing*

All behavioral testing was conducted during the active/light cycle under red light conditions. Behavioral testing was recorded (EthoVisionXT 13, Noldus) from top-view (Monochrome Gig-E Camera) at 30 frames/second and 600x800 pixels for later analysis.

**Social interaction test (SIT).** Mice were assessed on a two-trial social interaction test one day following CSW/DS (Fig. 1a). In the first 2.5-minute trial (No Target trial), the experimental mouse was allowed to freely explore the 44 cm x 44 cm white acrylic arena. A clear plexiglass and wire mesh enclosure (10 cm x 6 cm) was centered on one wall. In the second trial (Target trial), a novel CD-1 aggressor mouse was placed in the plexiglass and wire mesh enclosure and the experimental mouse was returned to the arena and allowed to freely explore for 2.5 minutes.

**Open field test (OFT).** Mice were individually placed into a corner of a white acrylic open arena (44 cm x 44 cm) and allowed to freely explore for 5 minutes.

*Calculation of motion and location characteristics*

To calculate total distance, center torso distance between consecutive frames is calculated and summed for each syllables' occurrence (i.e., total pixel distance). Syllable velocity is calculated from distance by the number of frames of a given syllable's occurrence (i.e., total pixel distance / n frames). Spine length is calculated as the average distance between front torso and tail base, whereas head length is calculated as the average distance between snout and front torso. Finally, distance to center and distance to target are the average distance of center torso to the respective points of interest.

*Frame Independent Projected Fiber Photometry*

To measure calcium-associated changes in fluorescence in real time, recordings were made from mPFC NAc-projecting cells during the social interaction following CSDS. Samples were collected at a frequency of 19.5 Hz using Neurophotometrics hardware through Bonsai and FlyCap software. Recordings were coupled to the start of behavioral analysis by interfacing Bonsai with MED-PC using a custom DAQ box (Neurophotometrics). Data were extracted and analyzed using custom-written scripts in Python and R (see code availability statement) To normalize the data and control for potential artifacts, the control channel (415nm) was fitted to the raw (470nm). The fitted control was then subtracted from the raw trace. The resultant trace was divided by the fitted control, giving the $\Delta F/F$. No target and target trials were concatenated and converted to a z-score for each animal.

### Supplemental Figures

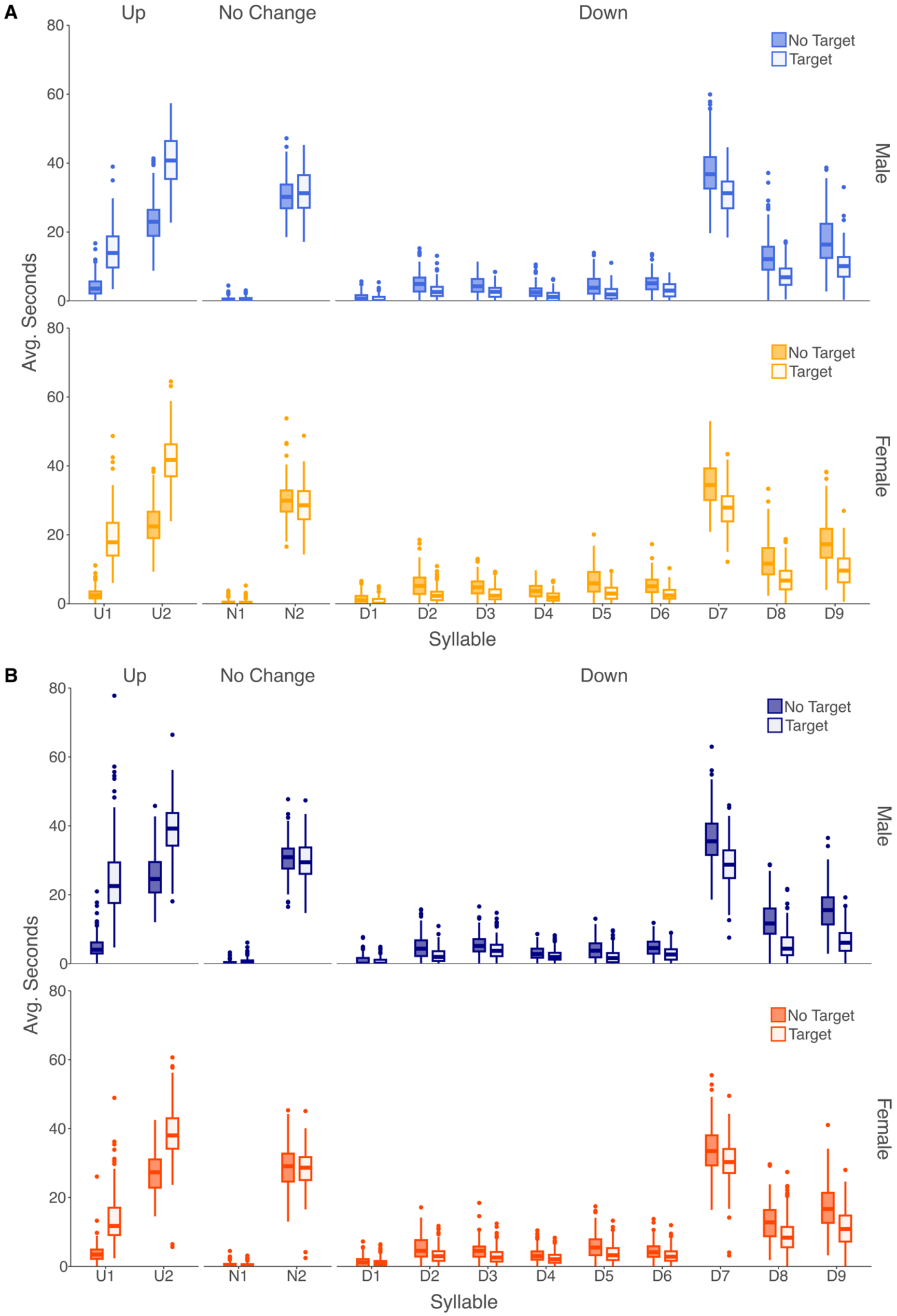


**Figure S1. Raw syllable frequencies by condition and sex.** (A) Raw syllable frequency in No Target (filled) and Target (empty) trials in control males (top panel) and females (bottom panel). (B) Raw syllable frequency in No Target (filled) and Target (empty) trials in stressed males (top panel) and females (bottom panel).

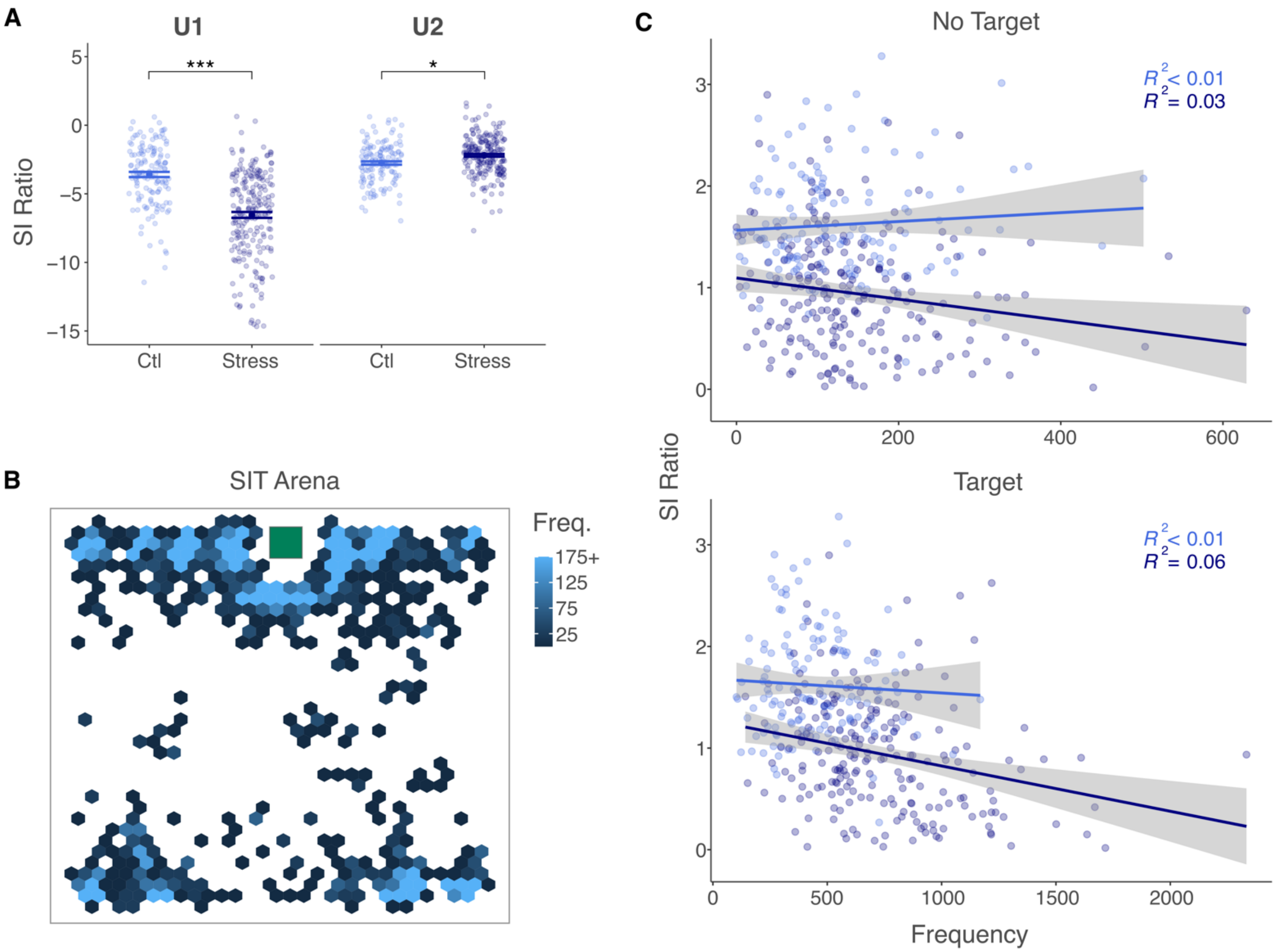


**Figure S2. Male stress-effects on Upregulated syllables are driven by U1.** (A) Stressed males display a higher upregulation of U1 and a marginally lower upregulation of U2 than controls (ANOVA; Condition: $F_{1,338} = 51.15$, $p < 0.001$; Syllable: $F_{1,355} = 304.6$, $p < 0.001$; Condition x Syllable: $F_{1,355} = 104.6$, $p < 0.001$; Cohort: $F_{17,338} = 3.196$, $p < 0.001$; Sidak Post-Hoc: Control U1 *vs* Stress U1: *t-ratio* = 12.148, $p_{adj} < 0.001$; Control U2 *vs* Stress U2: *t-ratio* = -2.328, $p_{adj} < 0.05$). (B) Syllable U1 is largely expressed around the mesh enclosure (green square) and in the corners in an example cohort (E15). (C) Frequency of U1 correlates with SI ratio in stressed males but not controls (Control NT: $r_{Pearson} = 0.074$, $p = 0.384$; Stress NT: $r_{Pearson} = -0.178$, $p < 0.01$; Control T: $r_{Pearson} = -0.052$, $p = 0.537$; Stress T: $r_{Pearson} = -0.254$, $p < 0.001$)

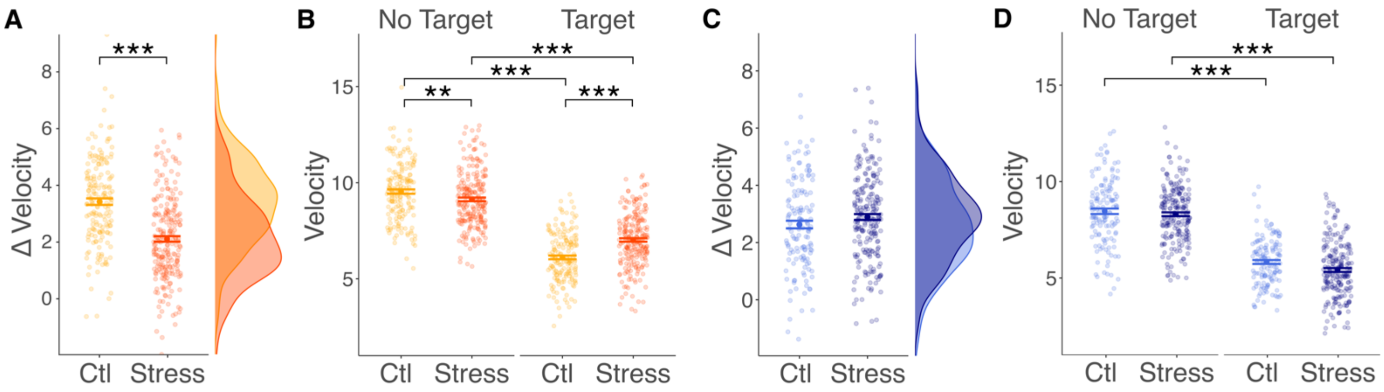


**Figure S3. Stress effects on ΔVelocity are female-specific and largely driven by increased velocity of stressed females in the Target trial.** ΔVelocity is significantly decreased in stressed females (A) but not stressed males (C) compared to controls (ANOVA; Condition: $F_{1,738}$ = 32.225, $p < 0.001$; Sex: $F_{1,738}$ = 1.72, $p = 0.19$; Condition x Sex: $F_{1,738}$ = 60.568, $p < 0.001$; Cohort: $F_{19,738}$ = 8.523, $p < 0.001$; Sidak Post-Hoc: Control Male *vs* Stress Male: *t-ratio* = -1.645, $p_{adj}$ = 0.1; Control Female *vs* Stress Female: *t-ratio* = 9.657, $p_{adj} < 0.001$). (B) While both control and stressed females decrease in Velocity in the Target trial compared to the No Target trial, stressed females display a lower velocity than controls in the No Target trial and a higher Velocity than controls in the Target trial, with a larger effect in the Target trial (repeated measures ANOVA; Condition: $F_{1,385}$ = 5.254, $p < 0.05$; Trial type: $F_{1,402}$ = 1295.13, $p < 0.001$; Condition x Trial type: $F_{1,402}$ = 76.56, $p < 0.001$; Cohort: $F_{17,385}$ = 6.108, $p < 0.001$; Sidak Post-Hoc: No Target Control *vs* No Target Stress: *t-ratio* = 3.206, $p_{adj} < 0.01$; Target Control *vs* Target Stress: *t-ratio* = -6.685, $p_{adj} < 0.001$; No Target Control *vs* Target Control: *t-ratio* = 30.22, $p_{adj} < 0.001$; No Target Stress *vs* Target Stress: *t-ratio* = 21.412, $p_{adj} < 0.001$). (D) In contrast, both control and stressed male mice show a decrease in Velocity from Target to No Target trial, with no differences between stress and control group (repeated measures ANOVA; Condition: $F_{1,338}$ = 5.856, $p < 0.05$; Trial type: $F_{1,355}$ = 1177.303, $p < 0.001$; Condition x Trial type: $F_{1,355}$ = 2.498, $p = 0.115$; Cohort: $F_{17,338}$ = 7.132, $p < 0.001$; Sidak Post-Hoc: No Target Control *vs* No Target Stress: *t-ratio* = 0.567, $p_{adj}$ = 0.571; Target Control *vs* Target Stress: *t-ratio* = 2.382, $p_{adj}$ = 2.382; No Target Control vs Target Control: *t-ratio* = 20.334, $p_{adj} < 0.001$; No Target Stress *vs* Target Stress: *t-ratio* = 27.683, $p_{adj} < 0.001$). *$p_{adj} < 0.05$ **$p_{adj} < 0.01$ ***$p_{adj} < 0.001$.

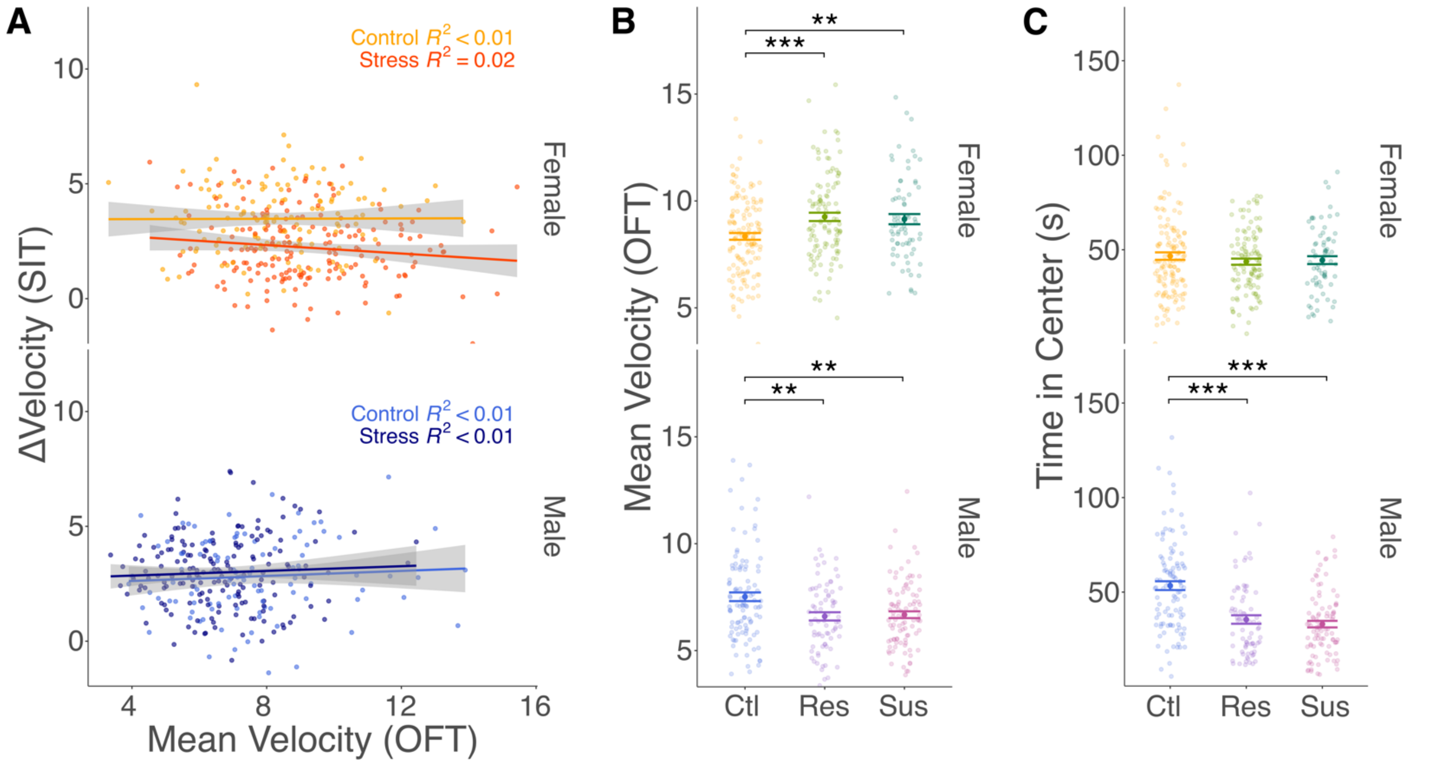


**Figure S4. Resilient and susceptible females show anxiety-like behavioral changes in Velocity in the OFT.** (A) ΔVelocity does not correlate with Velocity in the OFT, further confirming that ΔVelocity changes are driven by the presence of an aggressive target rather than underlying individual differences in locomotion (Females: Controls: $r_{Pearson} = 0.004$, $p = 0.959$; Stress: $r_{Pearson} = -0.124$, $p = 0.097$, Males: Controls: $r_{Pearson} = 0.069$, $p = 0.483$; Stress: $r_{Pearson} = 0.053$, $p = 0.489$). (B) Both susceptible and resilient females show increased Velocity in the OFT, suggesting an overall change in anxiety-like behaviors after chronic stress exposure. In contrast, both susceptible and resilient males show a decrease in Velocity in the OFT compared to controls (ANOVA; Sex: $F_{1,571} = 157.146$, $p < 0.001$; Group: $F_{1,571} = 2.675$, $p = 0.070$; Sex x Group: $F_{1,571} = 15.020$, $p < 0.001$; Cohort: $F_{15,571} = 6.190$, $p < 0.001$; Sidak Post-Hoc: Female Ctl *vs* Female Res: *t-ratio* = -3.780, $p_{adj} < 0.001$; Female Ctl *vs* Female Sus: *t-ratio* = -3.175, $p_{adj} < 0.01$; Female Res *vs* Female Sus: *t-ratio* = 0.161, $p_{adj} = 0.986$; Male Ctl *vs* Male Res: *t-ratio* = 2.961, $p_{adj} < 0.01$; Male Ctl *vs* Male Sus: *t-ratio* = 3.049, $p_{adj} < 0.01$; Male Res *vs* Male Sus: *t-ratio* = -0.114, $p_{adj} = 0.993$). (C) Stressed females do not show differences in the usually employed metric Time in Center, whereas both susceptible and resilient males spend less time in the center of the OFT compared to controls (ANOVA; Sex: $F_{1,571} = 3.532$, $p = 0.061$; Group: $F_{1,571} = 22.215$, $p < 0.001$; Sex x Group: $F_{1,571} = 12.141$, $p < 0.001$; Cohort: $F_{15,571} = 6.846$, $p < 0.001$; Sidak Post-Hoc: Female Ctl *vs* Female Res: *t-ratio* = 1.638, $p_{adj} = 0.231$; Female Ctl *vs* Female Sus: *t-ratio* = 0.346, $p_{adj} = 0.936$; Female Res *vs* Female Sus: *t-ratio* = -1.035, $p_{adj} = 0.555$; Male Ctl *vs* Male Res: *t-ratio* = 5.822, $p_{adj} < 0.001$; Male Ctl *vs* Male Sus: *t-ratio* = 6.957, $p_{adj} < 0.001$; Male Res *vs* Male Sus: *t-ratio* = 0.625, $p_{adj} = 0.807$). **$p_{adj} < 0.01$ ***$p_{adj} < 0.001$.
